# Evolutionary trends in *Bombella* apis CRISPR-Cas systems

**DOI:** 10.1101/2025.02.05.636693

**Authors:** Carrie L Ganote, Lílian Caesar, Danny W Rice, Rachel J Whitaker, Irene LG Newton

## Abstract

Bacteria and archaea employ a rudimentary immune system, CRISPR-Cas, to protect against foreign genetic elements such as bacteriophage. CRISPR-Cas systems are found in Bombella apis, a microbe associated with honey bee queens, brood, and royal jelly. Unlike other honey bee microbiome members, B. apis does not colonize the worker bee midgut or hindgut and has therefore been understudied with regards to its importance in the honey bee colony. However, B. apis appears to play beneficial roles in the colony, by protecting developing brood from fungal pathogens and by bolstering their development under nutritional stress. Previously we identified CRISPR-Cas systems as being acquired by B. apis in its transition to bee association, as they are absent in a sister clade. Here we assess the variation and distribution of CRISPR-Cas types across B. apis strains. We found multiple CRISPR-Cas types, some of which have multiple arrays, within the same B. apis genomes and also in the honey bee queen gut metagenomes. We analyzed the spacers between strains to identify the history of mobile element interaction for each B. apis strain. Finally, we predict interactions between viral sequences and CRISPR systems from different honey bee microbiome members. Our analyses show that the B. apis CRISPR-Cas systems are dynamic, that microbes in the same niche have unique spacers which supports the functionality of these CRISPR-Cas systems, and that acquisition of new spacers may be occurring in multiple locations in the genome, allowing for a flexible antiviral arsenal for the microbe.

**IMPORTANCE:** Honey bee worker gut microbes have been implicated in everything from protection from pathogens to breakdown of complex polysaccharides in the diet. However, there are multiple niches within a honey bee colony that host a different group of microbes, including the acetic acid bacterium *Bombella apis. B. apis* is found in the colony food stores, in association with brood, in worker hypopharyngeal glands, and in the queen digestive tract. The roles that *B. apis* may serve in these environments are just beginning to be discovered and include production of a potent antifungal that protects developing bees and supplementation of dietary lysine to young larvae, bolstering their nutrition. Niche specificity in *B. apis* may be affected by the pressures of bacteriophage and other mobile elements which may target different strains in each specific bee environment. Studying the interplay between *B. apis* and its mobile genetic elements (MGEs) may help us better understand microbial community dynamics within the colony and the potential ramifications for the honey bee host.

## INTRODUCTION

Honey bees are an amazing model system for studying the interplay between society and the microbiome. The honey bee is a massively successful superorganism that lives as a eusocial colony, with one reproductively capable member (the honey bee queen), who generates all the offspring. Effectively sterile female worker bees complete all required tasks in the colony from rearing the next generation, to tending the queen, to foraging for resources. As part of the collective, honey bees exchange gut contents with each other through a process called trophylaxis - their foregut (also called the crop) containing nectar or other nutrients is transferred to other adult bees and developing brood in the colony, disseminating the nutrition gathered from flowers and other nectaries. This phenomenon has led to the foregut of the bee being termed the “social stomach.” In this way, one can imagine that microbes (including phages) associated with these animals would be introduced to the colony through the social interaction between the insects after foraging.

### Acetic acid microbes show consistent association with honey bees and their environments

*Bombella apis* is an important acetic acid bacterium found in honey bee food stores, royal jelly, the queen bee digestive tract, the larval bee gut, and in the hypopharyngeal glands of nurse bees (1). Workers use these glands to produce royal jelly, which they feed to developing brood and queens. Interestingly, *B. apis* is the only honey bee associated microbe identified that can withstand the antimicrobial properties of royal jelly (2). *B.apis* is a gram-negative Alphaproteobacterium within the family *Acetobacteraceae* and generally considered a beneficial microbe to bees, as it has been shown to have an inhibitory effect on pathogenic fungi (3), and can bolster honey bee nutrition by supplementing lysine to developing larvae (2). Based on 16S rRNA gene sequencing, a single bee colony can harbor many strains of *B.apis* (4).

### Microbial adaptive immunity

In the environments that *B. apis* inhabits inside a hive, phage predation may play a role in shaping its ecology and evolution. One of the tools in the prokaryotic arsenal against viral threats is CRISPR (Clustered Regularly Interspersed Short Palindromic Repeats) and CRISPR-associated (Cas) proteins (5). In this system, snippets of DNA, called spacers, are stored in the genomes of many bacteria and archaea interspersed with repeats (6, 7, 8, 9). Spacers match sequences from bacteriophage, plasmids or other mobile genetic elements (MGEs) and confer resistance against them (10) by acting as guide RNA to degrade matching sequences (11, 12).

The diverse CRISPR-Cas systems are categorized into types that differ in protein function and in the sequence that makes up the direct repeat (12). For this study, we will focus on types I-E and II-C, as they are prevalent across *Bombella*. Previous research has provided a model for the structure of Type I-E CRISPR-Cas systems (13), characterized by a palindromic repeat that forms a hairpin structure. The hairpin structure constrains selection on the sequence; if a mutation were to occur in the stem of the hairpin, it could potentially disable it. The top is a 4-bp loop that does not appear constrained. In Type II-C systems, Cas9 is equipped with its guide RNA a little differently. It encodes a repeat as well as a reverse complement of its repeat (called the tracrRNA), which act together to load the target spacer (12). In both types, the guide CRISPR RNA (crRNA) is loaded onto Cas endonucleases to seek, cleave, and disarm matching sequences within the cell (14).

### Previous work

When *B. apis* genomes were first analyzed, CRISPR-Cas loci were identified as distinguishing this genus from its sister group Saccharibacter (15) and the systems were hypothesized to be active based on the diversity of CRISPR-Cas spacers in the genomes of sequenced isolates.

In this study, we explored the hypothesis that mobile elements, such as bacteriophage, are active in bee-associated *B. apis* by identifying and characterizing the CRISPR-Cas loci found across all sequenced strains in this group of *Acetobacteraceae*. In addition to previously published genomes, samples from honey bee queen metagenomes were analyzed to focus on dynamics within an individual organism.

## RESULTS

### *Bombella* genomes contain two CRISPR-Cas system types

We used the 16 available *Bombella* genomes to discover all CRISPR-Cas systems present in this clade. In *B. apis* and its close relatives, we see two CRISPR-Cas system types - type I-E and type II-C. Both of these system co-occur on the genome in most strains Figure 3. Models for the structure of CRISPR-Cas genes and arrays are shown in Figure 1. The hairpin structure typical in type I-E CRISPRs can be seen, along with the 4bp tetraloop sequence with variations highlighted in yellow. The different tetraloop variations are also shown as different columns in Figure 3. Single-nucleotide polymorphisms (SNPs) can be found in the tetraloop portion and in the tails on either side of the hairpin without hindering the functionality of the repeat. Type II-C arrays differ only in the first three bases across *B. apis* strains, diverging further among *Acetobacteraceae* in bacterium AAB2 880 (see table S1).

**FIG 1.**
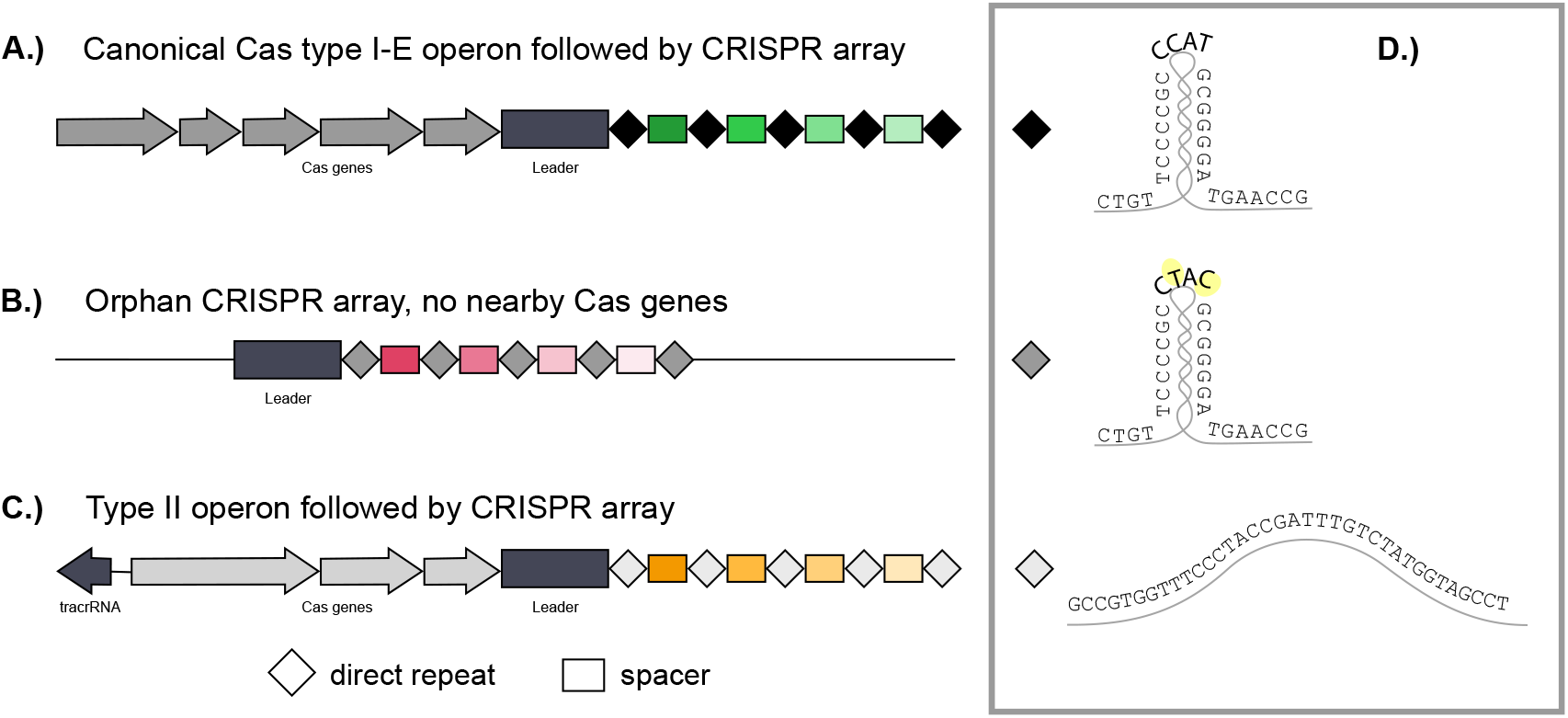
Schematic model for CRISPR Systems found in *B.apis* genomes. Type I-E and Type II-C CRISPR systems are both found in most *B.apis* genomes, with two arrays for type I-E. A.) Shows the typical layout for genes in the CRISPR system arranged as an operon, followed by a leader sequence and then the repeat-spacer array. B.) Show the secondary type I-E array prevalent in most *B.apis* genomes, called “orphan” here as no Cas genes are present nearby. C.) Shows the typical type II-C CRISPR system, including the tracrRNA and operon followed by the array. In D.), the structure of the direct repeat found in each array is shown. Nucleotides highlighted in yellow differ between canonical and orphan arrays. The sample type II-C repeat shown is from *B.apis* strain SME1.

### *Bombella apis* type I-E systems often use two distinct arrays

We further characterized the CRISPR-Cas systems in *Bombella* genomes to discover any patterns in the number of arrays present for each type. Analysis of the CRISPR-Cas systems reveals multiple Type I-E CRISPR arrays: one array in “canonical” formation, with the CRISPR array in close proximity (within 500bp) to the Cas operon associated with that CRISPR type, and the “orphan” array, far away from the protein-coding Cas genes that act upon it. In some cases, the exact distance between the array and associated Cas genes is unknown due to fragmented genome assembly. Long-read sequencing could be used to fully resolve relationships between CRISPR arrays by allowing for more complete, accurate resolution of these genomes.

In addition to being located away from the Cas operon, the orphan array harbors variations in the direct repeat tetraloop compared to the canonical array; Figure 1D highlights these differences in yellow. Orphan arrays have nearly double the number of spacers in their repertoire compared to canonical arrays (averages 30.9 and 18.9, resp.) Figure 2A. However, this difference is not significant (W = 76.5, p = 0.1084). When looking at the differences in number of spacers between type I-E and type II-C in *Bombella* genomes, there was a significant difference (W = 258, p = 0.0001) Figure 2B. Type II-C systems tend to have small arrays with a maximum of 12 and an average size of 5.9 spacers. This is compared to 25.9 on average for the I-E arrays (canonical and orphan taken together), with the largest array sporting 101 spacers (Figure 2).

**FIG 2.**
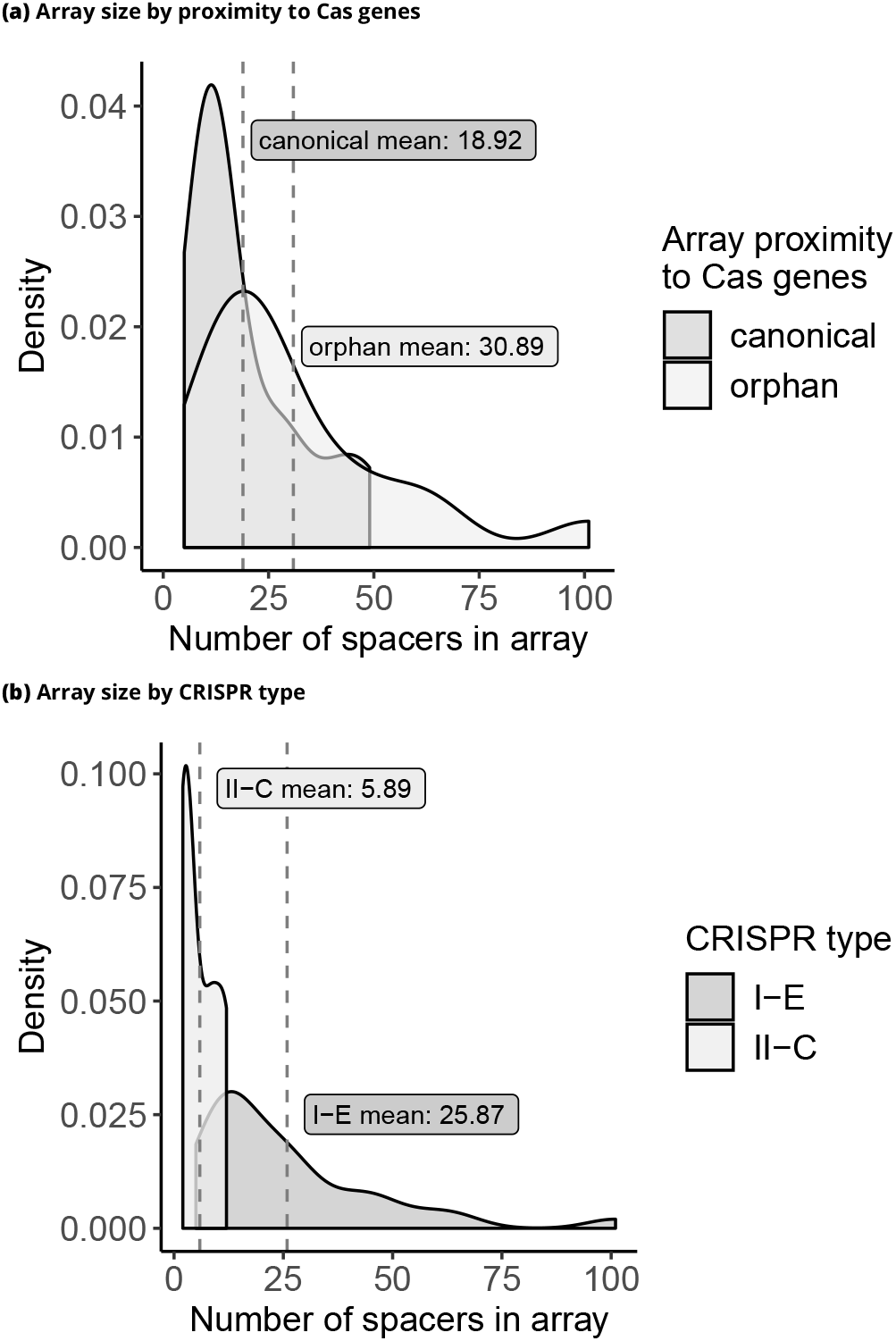
Array sizes differ among *Bombella* genomes. A.) Canonical CRISPR arrays immediately follow their associated Cas operon and tend to have fewer spacers than orphan arrays (no nearby Cas genes). B.) There is a difference in the average size of Type I-E CRISPR arrays and the smaller Type II-C CRISPR arrays in *Bombella* genomes.

**FIG 3.**
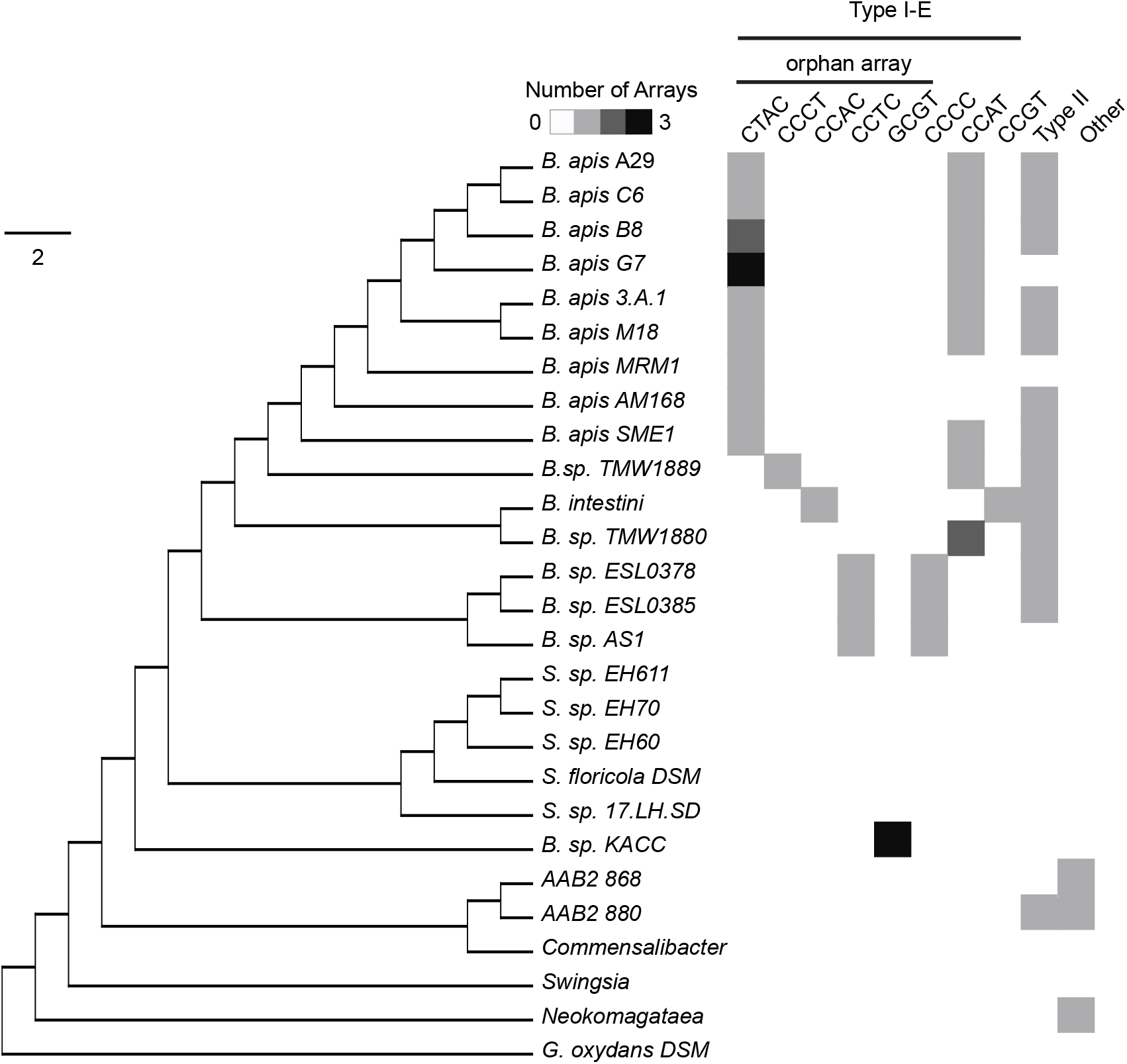
CRISPR Systems found across *Acetobacteraceae*. Cladogram depicting *Bombella apis* and related *Bombella* species with reference to other non-insect associated *Acetobacteraceae* such as *Swingsia, Neokomagataea*, and *Gluconobacter oxydans*. The tree was generated using the Gloome software with orthologs shared among these genomes. Columns indicate presence of different types of CRISPR arrays, with the number of arrays indicated by shades of gray, with darker color meaning more arrays. CRISPR arrays are further broken down for type I-E by the specific tetraloops in the repeats, as well as whether the array occurs as an orphan (no nearby Cas genes). The ‘Other’ column can include an array of any type other than those specified by other columns.

### *Bombella* CRISPR systems likely evolved within the clade and are mostly absent in related *Acetobacteraceae*

We used homology searches and ortholog clustering to identify the presence or absence of the above mentioned CRISPR systems across *Acetobacteraceae*. The genus *Saccharibacter* is a sister clade to the bee-associated *Bombella*, but thought to associate with pollen, floral nectaries or solitary bees (16)(17). Interestingly, although *Saccharibacter sp*. are closely related to *Bombella* species, they lack any intact CRISPR systems (Figure 3). One strain, *Saccharibacter sp. 17.LH.SD*, was found to contain a handful of type II Cas genes, but no array. As one moves further from *B. apis* in the phylogeny, it appears the type II-C system is not present reliably. For example, in *B. sp. KACC* (recently renamed *Aristophania vespae*), canonical CRISPR arrays were not found. Orphan arrays are placed far apart in this genome assembly and contain 4, 6, 4, and 3 spacers respectively. The type II-C system may have been acquired separately by the ancestor to *Bombella* and AAB2 880, as the repeat sequence is quite different in AAB2 880 compared to the rest of the type II-C systems found and it is absent in other members of this group.

### Few shared spacers across *B. apis* strains suggests active and dynamic systems

We hypothesize that if CRISPR systems are functional, they will actively recruit spacers and therefore spacer composition will differ between strains. We compared spacers found across *Bombella* genomes and found the vast majority of the spacers were unique to a single strain, suggesting that each strain has a separate history of phage and mobile genetic element interaction.

Spacers that were found in multiple genomes are shown in Figure 4 and in table S2 *B. apis* strain A29 shares one spacer with strain 3A1 and with *Bombella favorum*, and two spacers with *B. intestini. B. intestini* also shares two spacers with *B. apis* strain G7 and one spacer each with *Bombella* sp. ESL0378 and *Bombella* sp. ESL0385. Two shared spacers were identified between *B. apis* strains 3A1 and AM168, one spacer between SME1 and G7, one spacer between *Bombella* sp. AS1 and ESL0378. An anomalous 17 spacers are shared between ESL0378 and ESL0385, but these two genomes have the highest number of spacers in their arrays in general. While these cases show that there is some overlap between the spacers among the *Bombella* genomes, the vast majority of spacers were unique to their own genome. This abundance of unique spacers, even among closely related strains, suggests an active CRISPR system in *B. apis*.

**FIG 4.**
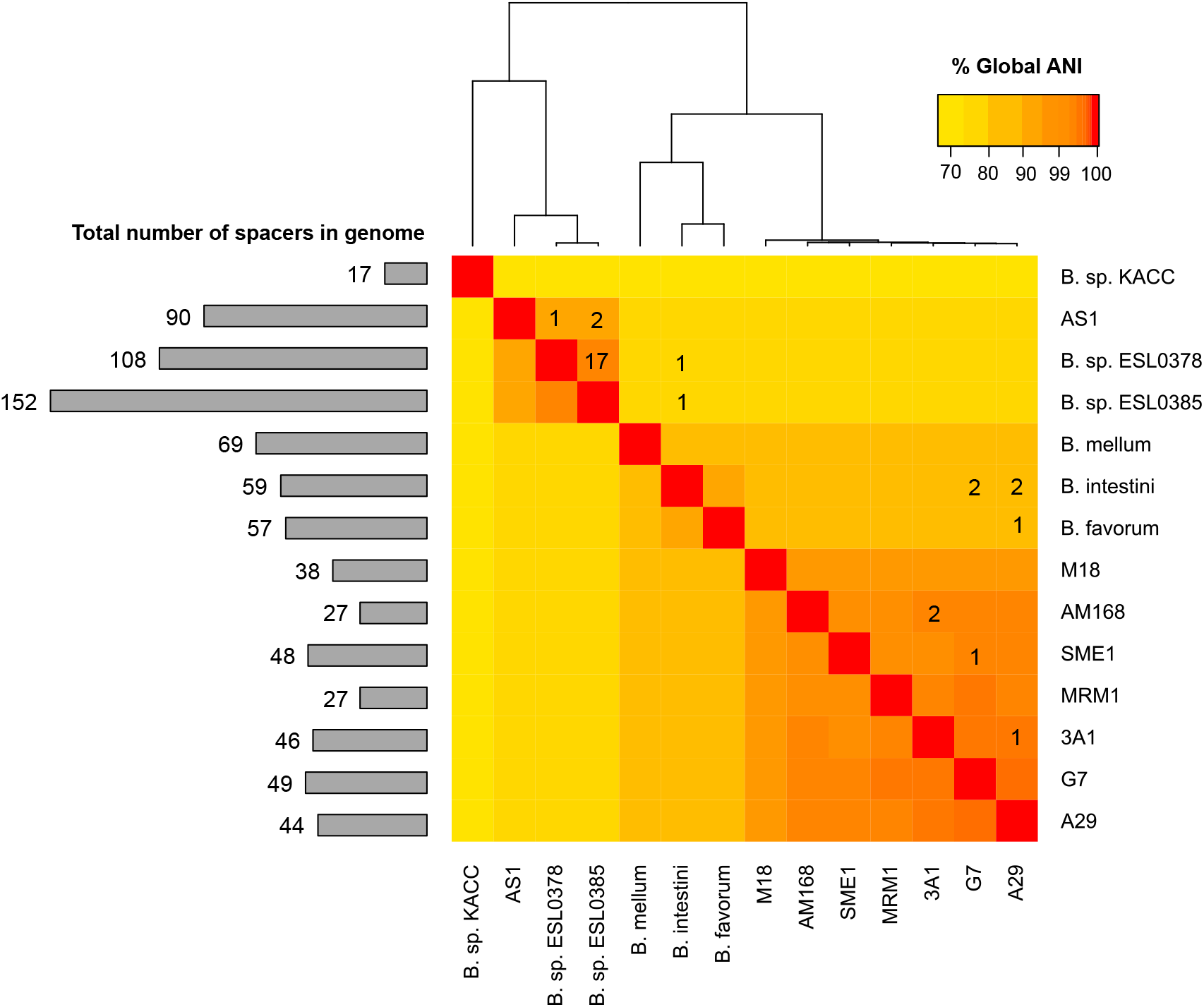
*Bombella* strains do not share many CRISPR spacers, despite genome-wide identity. Numbers shown in the heatmap indicate number of spacers with at least 80% sequence homology. Color of the map indicates the average nucleotide identity (ANI) between genomes. Note the color scale is skewed to better visually resolve *B. apis* strains, which all share over 99% identity. Cladogram representation of ANI is displayed on top. To the left of the heatmap is a histogram depicting the total number of spacers in each genome.

### Spacers from predicted CRISPR reads in queen metagenomes match spacers found in *Bombella* genomes

We explored metagenomic samples from queen bee guts to determine the prevalence of CRISPR constructs. Previous work on these data detected *Bombella* in ten queen metagenome samples (18). Six samples contained at least one predicted spacer that matched a spacer in *Bombella* genomes (D5, H4, E5, G8, A7, C8). *Bombella* sp. AS1, ESL0378, and ESL0385 each shared at least one spacer with one sample; *B. intestini, B. apis* 3A1 and A29 each shared at least one spacer with one sample, and strain SME1 shared at least one spacer with two different queen metagenome samples. *B. favorum* and *B. apis* G7 each shared at least one spacer with three different samples. Importantly, these metagenomes contain entire communities and not only *B. apis*. However, homology of the repeats associated with these spacers does suggest that these are CRISPR spacers from *B. apis*. More interesting is the fact that the isolates from published genomes and the CRISPR spacers we predicted in these metagenome samples may have shared MGE history in spite of different temporal, geographic, and niche sampling.

### Spacer sequences from *Bombella* CRISPR arrays match against prophage found in *Bombella* genomes

We compared spacers found in CRISPR arrays in *Bombella* genomes against prophage regions that were predicted in several *B. apis* strains(15) to test for evidence of active defense against these viruses. We used Phaster (19),(20) to identify regions in the *Bombella* genomes that are predicted to have phage origin. Ten spacers from type II-C CRISPR arrays matched against prophage regions, compared with 139 spacers from type I-E CRISPRS.

### Spacers from honey bee queen samples match prophage regions

To find evidence of prophage from published genomes triggering spacer acquisition in individual bee microbiomes, we tested for matches to spacers from honey bee queen samples (labeled in Figure 5 by a letter/number combination). Samples vary dramatically in which and how many spacers match prophage regions predicted in published genomes. Because of the nature of short-read metagenome assemblies, single-chromosome assemblies of *B. apis* could not be resolved and the CRISPR arrays were instead predicted from read data alone. Matches to a *Bombella* prophage, from a spacer flanked by repeats matching those found in other *Bombella* CRISPR arrays imply that these arrays were from *B.apis* or a close relative. Sample E5 is particularly enriched in spacers that hit the intact prophage in SME1, but many connections can be seen between queen samples and incomplete prophage across *Bombella*.

**FIG 5.**
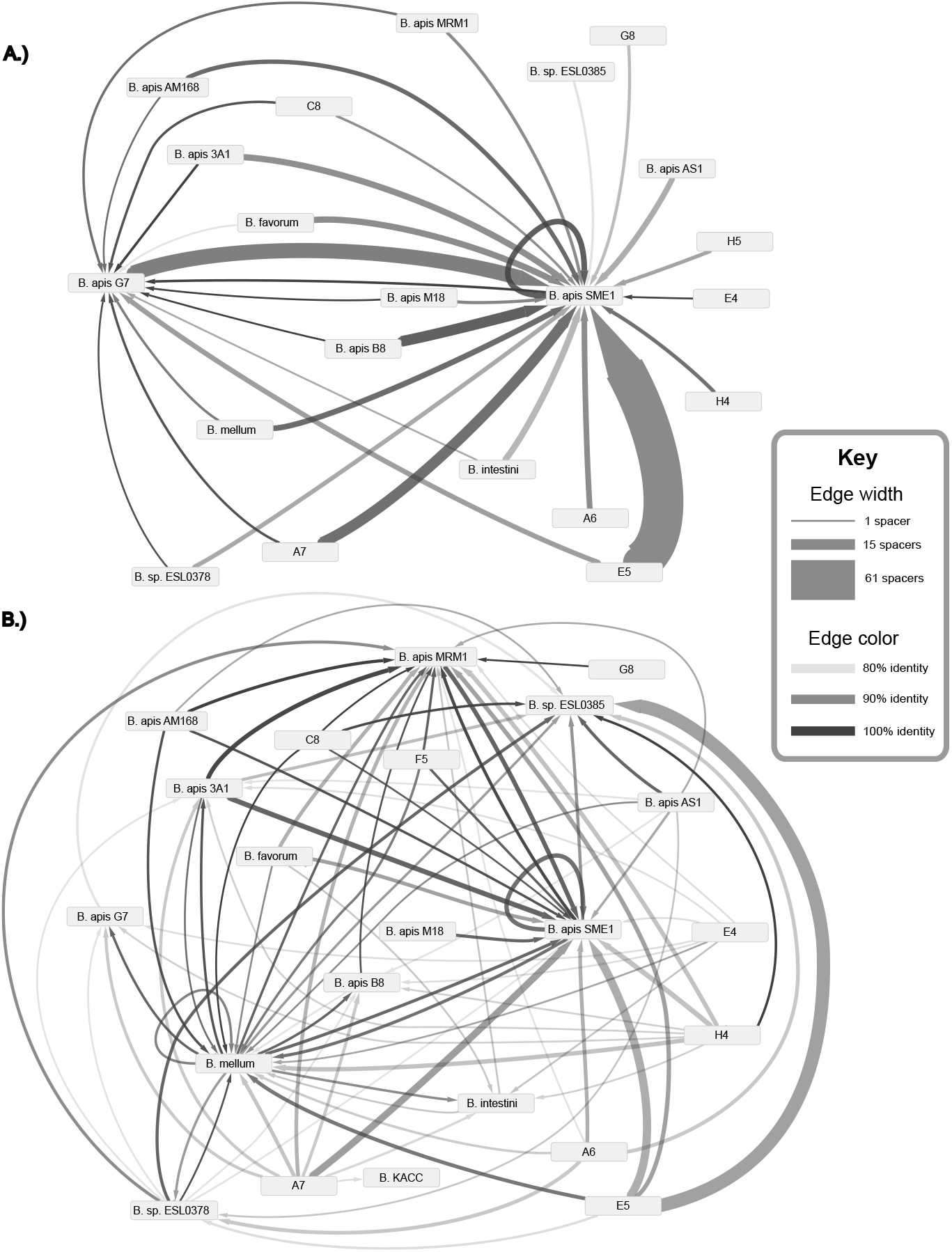
Matches between spacer sequences and prophage regions. Arrows point toward the genome containing the predicted prophage. The thickness of the lines depict how many spacer matches were found, while the color of the lines shows the average similarity in %matching nucleotides. A.) Intact prophage regions predicted by Phaster. B.) Incomplete/questionable prophage regions predicted by Phaster.

**FIG 6.**
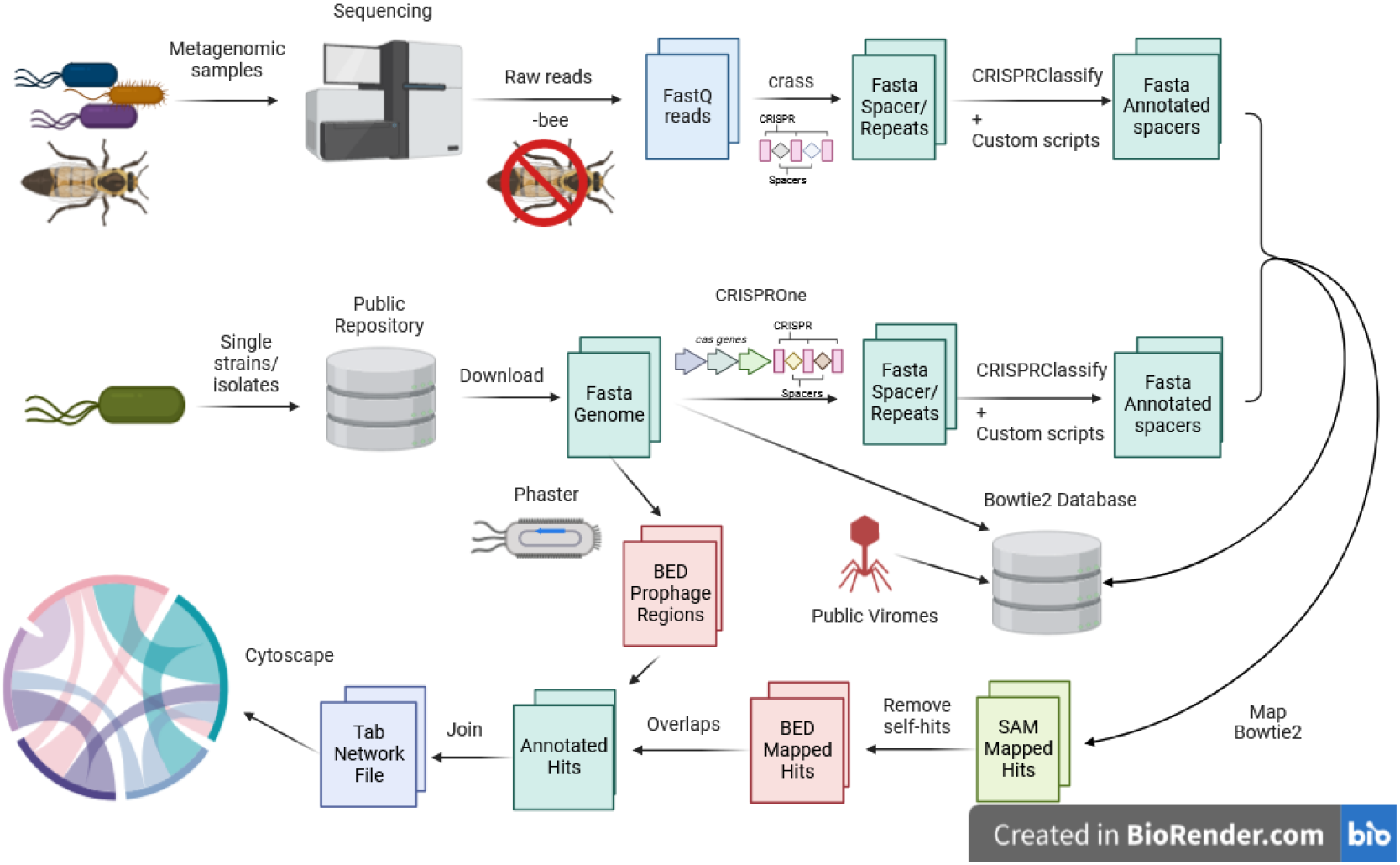
Bioinformatics Worklow schematic. **Genomes were sourced from public repositories and analyzed using CRISPROne to predict CRISPR regions. Metagenomic reads were analyzed with crass to pull out spacers and repeats. These spacers were mapped to known viruses from previous bee studies and to the genomes and other spacers; after cleaning up, the matches were visualized in Cytoscape.**

### Spacers from both CRISPR-Cas system types match against prophage regions of genomes

Matches between *Bombella* CRISPR spacers and putative prophage can be found from both the type I-E and the type II-C arrays, but they differ in number. Thirty spacers from type II-C systems matched against prophage regions, vs. 501 spacers from type I-E systems. Type II-C matches to the prophage regions may seem more rare, but the type II-C arrays in general are smaller than those of type I-E in *Bombella* genomes (Figure 2) and thus have fewer opportunities for spacer correlations in general. In no case did the same spacer sequence appear in both I-E and II-C systems; even if they matched against the same prophage, they picked up different protospacers from that phage. We saw no evidence that the same spacer sequence is occurring in both systems within the same genome.

### Spacer matches to genic regions

In addition to matches against the prophage regions in published genomes, 44 unique spacers matched to other parts of the genome. As *Bombella* strains are closely related (Figure 4), many of these spacers matched to multiple genomes, for a total of 187 matches. Of the 44 spacers that matched non-phage regions, 32 unique spacers landed within gene boundaries for a total of 142 matches.

### No spacer matches to viral bins or published viromes

Viral sequences from bacteriophage isolated from the worker gut (21), as well as predicted phage from the queen metagenome used in this analysis and in (18), were tested for spacer matches. However, no matches were found for *Bombella* species against any of these published viral sequences. This may reflect the lack of described phages for *Bombella* in the databases.

## DISCUSSION

Here we report a comprehensive analysis of CRISPR systems across the *Bombella* genus. This important microbe, found in honey bee association, has active CRISPR arrays and harbors prophages, suggesting an active arms race between the bacterium and its bacteriophages and other MGEs. Across *Bombella* genomes, CRISPR systems are conserved but not found consistently across more distantly related *Acetobacteraceae*; this may suggest that this defense mechanism was gained during association with honey bees (15, 2).

*B. apis* harbors two types of CRISPR systems - type I-E and type II-C. Utilizing a variety of CRISPR types may be beneficial to the bacteria if the systems differ in cleaving efficiency, as has been shown between types I-B and II-B (22). Additionally, most *B. apis* strains usually have two or more type I-E arrays, with one located right after the Cas genes and others far from them (usually on a different contig in the assembly); see Figure 3. Spacers in the leader end of the array are more abundantly transcribed into crRNA (22, 23); with more than one array, the bacterium could have more flexibility and speed in transcription. Better resolved *Bombella* genomes would more confidently place the location and orientation of the different CRISPR arrays.

Based on the different number of spacers as well as sequence identity between spacers across strains, we assert that the *Bombella* CRISPR systems are active. Our analysis of CRISPR homology to characterize past interactions suggests that many of these spacers target past lytic events of prophage infections. The difference between the number and strength of matches to the intact vs. incomplete/questionable prophage, as well as the higher prevalence of matches across longer evolutionary distance, suggests two hypotheses: (1) the spacer is an ancient relic from a prophage infection shared by a common ancestor, and time has mutated the spacer or the prophage since; (2) that it is a recent spacer acquisition of a phage that has mutated in the time since its predecessor integrated into the genome and degraded there.

There can be other explanations for differing spacers in CRISPR arrays other than Cas-mediated spacer acquisition. Spacers can be lost through deletions (24) and gained/lost through recombination events (25). There is a bias as to which spacers tend to be lost, with the tail end of the array usually more conserved than the middle or leader end (26). This is likely occurring in the samples we analyzed, but deletion alone doesn’t account for the unique spacers in each array. Recombination events seem to be more rare than deletion events reported in previous studies (27), and we do not find evidence of mixed repeats between arrays found in other studies (25).

Taken together, this paints a picture of spacer loss and replacement as expected in Cas systems, with newly acquired spacers replacing older ones. The relative lack of shared spacers among *Bombella* genomes implies that the CRISPR systems in these strains are working as intended, accumulating new targets for defense in dynamic interplay with bacteriophage and other mobile genetic elements. One curious result of our interrogation of the type I-E CRISPR arrays in *B. apis* is the discovery of multiple SNPs in the hairpin loop that is part of the CRISPR direct repeat. Under the assumption that selection on the loop portion of the direct repeat is neutral, the signature of drift in type I-E CRISPR repeats can be used as evidence that these CRISPR systems have been present and evolving in *Bombella* species for some time.

As of now, no *Bombella* phages have been isolated. Lytic phage from queen bee metagenomes were predicted to predate upon *Lactobacillaceae* and *Enterobacteriaceae*, but direct evidence linking phage to *Bombella* CRISPR spacers were not found (18). We were similarly unable to recover spacers matching lytic phage in the queen metagenomes, both from the same dataset (using crass instead of CRISPRFinder to call spacers) and from published genomes. Importantly, bacteriophage are not the only MGEs that CRISPRs block (24) - plasmids and transposons can also be the target of CRISPR-Cas immunity, and further studies should investigate other mobile elements within *Bombella* species.

Future experimental work could illuminate the mechanics between the two CRISPR system types in *B. apis*. If challenged with a phage known to target it, which CRISPR array will add spacers? More spacers are found in orphan arrays than canonical ones. What accounts for this difference - does the orphan array pick up spacers at a higher rate, or is it less likely to prune away the older spacers?

Honey bees acquire many microbes during their journeys outside and within the colony. MGEs are brought together and given a chance to associate inside the bee gut and food stores, increasing possible sources of genetic diversity for the microbes but also enabling hostile DNA to be acquired. Balancing these risks and opportunities may drive high turnover in CRISPR spacers.

## Conclusion

In conclusion, we explored the hypothesis that mobile genetic elements, such as bacteriophage, are actively eliciting an immune response in *Bombella apis* by characterizing the CRISPR-Cas loci found across all sequenced strains in this group of *Acetobacteraceae* as well as in metagenomes from queen bees. There is a strong signal of conservation in the CRISPR types and organization in *B. apis*. This conservation breaks down more distantly in the *Acetobacteraceae* tree among microbes not known for this specialized bee-associated niche. Even among closely related strains of *B. apis*, shared spacers are not common, supporting claims that these arrays are actively gaining and losing spacers. Spacers matching prophage sequences found in *B. apis* genomes provides evidence that bacteriophage are being targeted by CRISPR Cas systems in these microbes. These observations lead us to conclude that CRISPR systems in *B. apis* are actively responding immune systems against bacteriophage.

## MATERIALS AND METHODS

See 6 for a graphical depiction of the bioinformatics workflow used to generate CRISPR relationships.

### Data sources

Publicly available genomes were collected from NCBI for published strains of *Bombella apis* as well as related *Acetobacteraceae*, used as outgroups. See table S1 for full accession details. In addition, DNA sequences from the digestive tracts of ten queen honey bees from Vancouver apiaries (18) were used to explore CRISPR systems in *Bombella apis* as they exist together with any potential phage or niche-mates in the bee host. See table S2 for queen sample IDs and metadata.

### Phylogeny Estimation

RAxML (28) was used with the following options: -m GTRGAMMA -f a -N 100. A concatenated amino acid derived codon alignment was used containing 1,667,688 positions after gappy positions were removed. Ortholog codon alignments represented by 15 or more taxa were included in the alignment.

### Identifying and Categorizing Cas systems in B. apis strains

Putative CRISPR-Cas systems and their subtypes were predicted from the published genomes by running CRISPROne (29). CRISPROne predicts CRISPR constructs first using CRISPR Recognition Tool (CRT) (30) and applies further quality control to avoid mislabeling any repetitive sequences as CRISPRs. Polished results are visualized as graphical schematics showing the relationship between Cas operons and CRISPR arrays in the genome. While this software is primarily intended for interactive use from a web front end, the software can also be run as a local script for greater scaling but without graphical output.

CRISPROne was run both from the command line and from the web browser to predict potential CRISPR arrays and Cas proteins. CRISPROne offers GFF formatted outputs to describe the annotation of Cas genes and CRISPR arrays, as well as the raw output from CRT. Spacers and repeats were collected in bulk from both of these outputs using in-house scripts available on Github (31). In the case of *Saccharibacter sp. 17.LH.SD*, an additional Blastp search was run on the Blast web server using default settings to identify predicted type II Cas genes.

CRISPRClassify (32) was run on the predicted direct repeat sequences to place them into respective CRISPR types and subtypes. CRISPRClassify predicts which CRISPR type and subtype a repeat may belong to, providing both an edit distance from a matching repeat in its database, as well as a probability score for the match according to its machine learning approach. For a repeat to be considered a true CRISPR repeat, it should score greater than 75% according to CRISPRClassify or have 4 or fewer edits between it and a known repeat in CRISPRClassify’s database. There are some edge cases where CRISPRClassify gave a very low score to a repeat that had only 2 mismatches and otherwise perfectly followed the pattern of its CRISPR type and subtype, thus score alone was too conservative of a measure for filtering non-CRISPR repeats. The annotated direct repeats were compared across strains. The presence of

CRISPR-Cas type II systems was noted for presence/absence among the genomes. CRISPR-Cas type I-E repeats were further analysed for two aspects - their exact direct repeat sequence, and their proximity to their Cas operon. Proximity was determined by comparing each CRISPR array’s coordinates to the closest predicted Cas genes in the gff output from command line results. Bedtools closest tool was used to calculate this distance. CRISPR arrays that were located on a different contig or more than 500bp away from and Cas gene was labeled as “orphans”. CRISPR arrays in tandem with these genes were labeled as “canonical”.

The exact direct repeat sequence of CRISPR-Cas type I-E sequences is constrained by the palindromic structure of the repeat, which must form a small hairpin when transcribed in order to properly arm the endonuclease with the spacer to form a mature CRISPR-RNA (12). As such, nucleotide differences between direct repeats tend to occur on the tails or loop of the hairpin. We focused on the 4-bp loop sequence, as it was sufficient to differentiate the direct repeats from one another.

### Analysis of CRISPR spacers

Spacers were collected from all the CRISPR arrays found in the public genomes. Command line utilities were used to count spacers, and custom scripts extracted exact matches between spacers across different strains (31). Spacer counts per array were tested for significant differences using Mann-Whitney-Wilcoxon tests, first comparing array types (IE vs. II), and then orphan vs. canonical type I-E. In some genomes, there were multiple copies of the same spacer internally - these were removed before counting cross-genome matches. In case of strand differences in the assemblies, exact matches of reverse complements were also included in the matches between strains.

### Identifying CRISPR spacers in honey bee queen metagenomes

To exclude false CRISPRs that might come from the honey bee host, raw reads mapping to honey bee were first stripped from the dataset. While trimming the raw data is usually recommended for any bioinformatics pipeline, our analysis is sensitive to false negatives in early steps. To preserve any possible match to reads containing CRISPR array sequences, we opted to map the raw data without trimming first.

Bowtie2-build (33) was used to create an index from the *Apis mellifera* assembly Amel_HAv3.1 (RefSeq accession GCF003254395.2). Raw reads from queen metagenomic samples were mapped against this database using bowtie2 with default parameters. To exclude reads that matched the honey bee host, the mapped results were given to samtools with a filter flag -f12, which removes read pairs that both map to the host. Samtools fastq was run on the filtered reads to get them back into fastq format, discarding supplementary and secondary mapped reads. Our dataset contained no singletons, so we set the -s flag for samtools fastq to avoid creating empty files.

We used the crass software (34) (version 1.0.1) to identify and assemble CRISPR arrays from the filtered raw reads. Crass searches reads for the presence of multiple instances of a 23-mer that are at least 26bp apart by default; it then undergoes refinement steps to increase confidence in final CRISPR calls. As crass ignores paired read information, left and right reads were given as separate inputs and created separate outputs. Crass was run with the default minimum number of repeats of 2 and also a cutoff of 2 for the minimum number of spacers to be considered as one group. Results were consolidated using crisprtools merge (part of the crass software family) and all direct repeats were separated into a file using crisprtools extract. CRISPRSClassify was run on the list of direct repeats in order to filter out repeats that do not belong to know CRISPR-Cas families.

Once the repeat type/subtype was classified and validated by CRISPRClassify, this information was propagated back to the consolidated results from crisprtools using custom scripts. This produced a list of all spacers found, along with information about which sample they originated from, what repeat and type they are associated with, and their reverse complement. Even with lenient parameters, crass failed to recover all reads that contained type I-E CRISPR sequences that are found in the *Bombella* clade; these were recovered from the raw reads using a combination of grep, custom scripts (31) and manual cleanup to ensure that these spacers were also included in the analysis of spacers in the metagenome. Since spacers were derived from raw reads rather than assemblies, it was possible to recover partial spacers at the front or end of a read; any partial spacer less than 16 bp long was discarded as this would lead to spurious mapping hits.

### Finding matches between honey bee queen metagenome CRISPR spacers and *Bombella* genomes

A reference sequence collection was built with published *Bombella* genomes, putative viral bins from the queen metagenome, and honey bee worker gut viromes found in (21). Bins from the queen metagenome were analyzed in (18), and a list of bins were found that are predicted to be lytic phage - see table S5 for a list of viral bins. Sequencing data of the two worker gut viromes were acquired from NCBI under BioProject PRJNA599270. The genomes of the eight pure phage isolates were acquired from GenBank, under accession numbers MT006233–MT006240. Additionally, the metagenome from the same study (GenBank: JAAOBB000000000.1) was included. This reference was indexed with bowtie2-build and all collected spacers were mapped using bowtie2 with the -a flag to report all matches and –no-unal to ignore spacers which did not align to any source in the reference. This produced a set of mapped spacers that allowed for some mismatch; in order to retrieve spacer matches with more lenient homology (down to 80% similarity), bowtie2 was run with the following parameters: -a –no-unal -D 20 -R 3 -L 11 -N 1 –gbar 1 –mp 3 -p 6. Mapped spacers were then filtered using custom scripts to remove self-hits; since all spacers were included in both the reference and the query set, self matches were guaranteed in the results. Spacer matches with identity less than 80% were discarded.

### Calling putative prophage regions in *Bombella* genomes

Phaster (19),(20) is a web-based prediction tool for finding evidence of phage sequence within a larger sequence input. Phaster was utilized to predict putative prophage regions within *Bombella* genomes given as input. In addition to general regions, individual phage proteins are annotated and delimited. Based on the presence of predicted proteins necessary for full phage viability, putative prophage regions are scored by Phaster as intact, questionable, or incomplete. For the purposes of this study, all phage regions were considered regardless of score.

Putative prophage regions were converted into intervals and output in BED format using custom scripts. Spacers which mapped to published *Bombella* genomes were similarly converted into intervals from the sam file into a bed file, however, 32bp of padding was added to the each side of the location reported by the sam file to allow some buffer in case of poor annotation boundaries and to account for any error in strand associated with the spacer. Bedtools intersect (35) was used with the -c flag to count the instances of overlap between spacers and each prophage region in each *Bombella* genome. Finally, overlaps were summed across prophage regions to get a count of spacers possibly derived from prophage sequence per genome. The %identity was calculated for each set of matches between a sample/genome and a prophage region, with incomplete and questionable being lumped together. Spacers which did not map within the prophage region, but did map within the genome, were found using bedtools subtract against the prophage regions and the CRISPR arrays themselves to avoid counting self-hits. These matches were then intersected with gene regions as pulled from GFF files corresponding to the genomes in order to find matches to genic regions annotated in these genomes.

### Visualization

Hits between spacers from different samples were pulled from the alignment and placed into a tab-delimited file for consumption by the network visualization software Cytoscape (36). This file contains the name of the query, the name of the reference, and how many spacers were matched.

Alignments to prophage regions were filtered from the alignment file and placed into a tab-delimited file containing the following: name of the query; name of the reference; whether the prophage region was intact or incomplete/questionable; the count for each of these categories, and the average %identity of the matches. Fed into Cytoscape, a directed graph was created and two networks are shown in Figure 5. In the first graph, nodes were filtered to only show connections to intact prophage (called complete in this study). The second graph illustrates incomplete prophages, which includes questionable and incomplete regions as predicted by Phaster.

Global average nucleotide identity (gANI) was calculated from CDS sequence files corresponding to each published Bombella genome using ANIcalculator v1 (37). Spacer-spacer matches were visualized in R using gANI to create a symmetrical heatmap (31). The numbers of shared spacers were added manually.

### Availability of data and materials

All data used in this study is available to the scientific community in the form of accession numbers for genomes used **??**, queen bee metagenome sets available from the publication (18), and in the code used to generate the results in this study (31).

## Data citation

## SUPPLEMENTAL MATERIAL

**TABLE S1**. Type II-C repeats identified across Bombella genomes.

**TABLE S2**. Number of shared spacers, gANI, and total mismatches across Bombella genomes.

**TABLE S3**. Accession numbers for the publicly available genomes used in this study.

**TABLE S4**. Queen metagenome metadata used in this study and previously published.

**TABLE S5**. ID of putative viral bins from metagenomic assemblies of the queen microbiome.

## ACKNOWLEDGMENTS

This work was supported by National Science Foundation IOS award 2005306 DBI Biology Integration Institutes Program, Award 2022049 (to ILGN). The authors acknowledge the Indiana University Pervasive Technology Institute for providing supercomputing and data storage resources that have contributed to the research results reported within this paper. (38)

